# How to quantify immigration from community abundance data using the Neutral Community Model

**DOI:** 10.1101/2025.04.12.648546

**Authors:** Ramis Rafay, Eric W. Jones, David A. Sivak, S. Jane Fowler

**Author notes:** **Corresponding author:** S. Jane Fowler, **Email:**. **Competing Interest Statement:** N/A.

## Abstract

Biological communities are connected through dispersal, which regulates diversity across local and regional scales. However, dispersal is difficult to measure directly, limiting what is known about dispersal’s impact on species composition in complex communities. One method to measure dispersal employs the Neutral Community Model (NCM) to quantify how a local community is influenced by the immigration of individuals from a larger source community. Conveniently, the immigration rate *N*_T_*m* of the NCM can be fit from biological sequence abundance datasets, which are plentiful. Yet it is neither known if these estimated values reflect the ground truth, nor what sampling effort is required to yield accurate estimates. In this study we introduce two inference methods, a variance-based and a Dirichlet-Multinomial Log-Likelihood (DM-LL) method, to complement the established occupancy-based inference method. In simulations of communities that resemble activated sludge microbiomes, all inference methods were capable of estimating *N*_T_*m* within 10% of ground-truth, with the variance-based and DM-LL methods requiring less sampling effort. Accurate inferences require read depths greater than *N*_T_*m* in each sample. The three methods agree in their inferred *N*_T_*m* in simulations of communities experiencing weak non-neutral effects (e.g., selection), and in applications to an empirical dataset from wastewater activated sludge. Based on these findings, we propose practical sampling and methodological guidelines for quantifying immigration between highly diverse, complex communities using the NCM.

## Introduction

A central goal in ecology is to describe how biological communities assemble and vary over time and space. The four fundamental community assembly processes—selection, speciation, dispersal, and drift—determine taxa composition and diversity (*i.e*., community structure) in any habitat (1). While selection, speciation, and drift operate at local scales, dispersal is a strictly regional process that connects communities across larger spatial scales through immigration and emigration. Moreover, the rate of dispersal mediates the relative influence of stochastic and deterministic assembly processes: high dispersal homogenizes communities and reduces the influence of selection and drift, while low dispersal increases the impact of local differentiation and taxa turnover on community structure (2). Advances in high-throughput sequencing methods enable researchers to census micro- and macro-organismal communities with unprecedented resolution (3, 4), facilitating the development of models that describe community assembly and community structure at finer spatial and temporal scales. The Neutral Community Model (NCM) has emerged as a powerful and analytically tractable framework for quantifying how dispersal affects community structure in ideal ‘neutral’ communities in which all taxa are assumed to have equal growth and death rates (5). Though biological communities are obviously not made of species with identical fitness, NCMs nonetheless recapitulate observed ecological patterns like taxa-area, and distance-decay relationships (6–9).

In the NCM, the abundance of each taxon in a ‘local’ community is determined by stochastic birth-death processes and immigration from an external ‘source’ community. At each timestep a random individual in the local community is lost to death or emigration, then replaced by (i) an immigrant from a larger source community with a probability *m*, or (ii) by the reproduction of a random individual in the local community with a probability *1*−*m* (Fig. 1). The long-term taxa abundance distributions predicted by the NCM are approximated by a Dirichlet distribution that depends on the immigration rate *N*_T_*m*, the expected number of immigrants colonizing the local community per *N*_*T*_ replacement events (10, 11). The larger the immigration rate, the more local-community composition is driven by immigration from external sources, which can be an important process for maintaining the presence of rare (low abundance) taxa that would otherwise go extinct due to drift.

**Figure 1.**
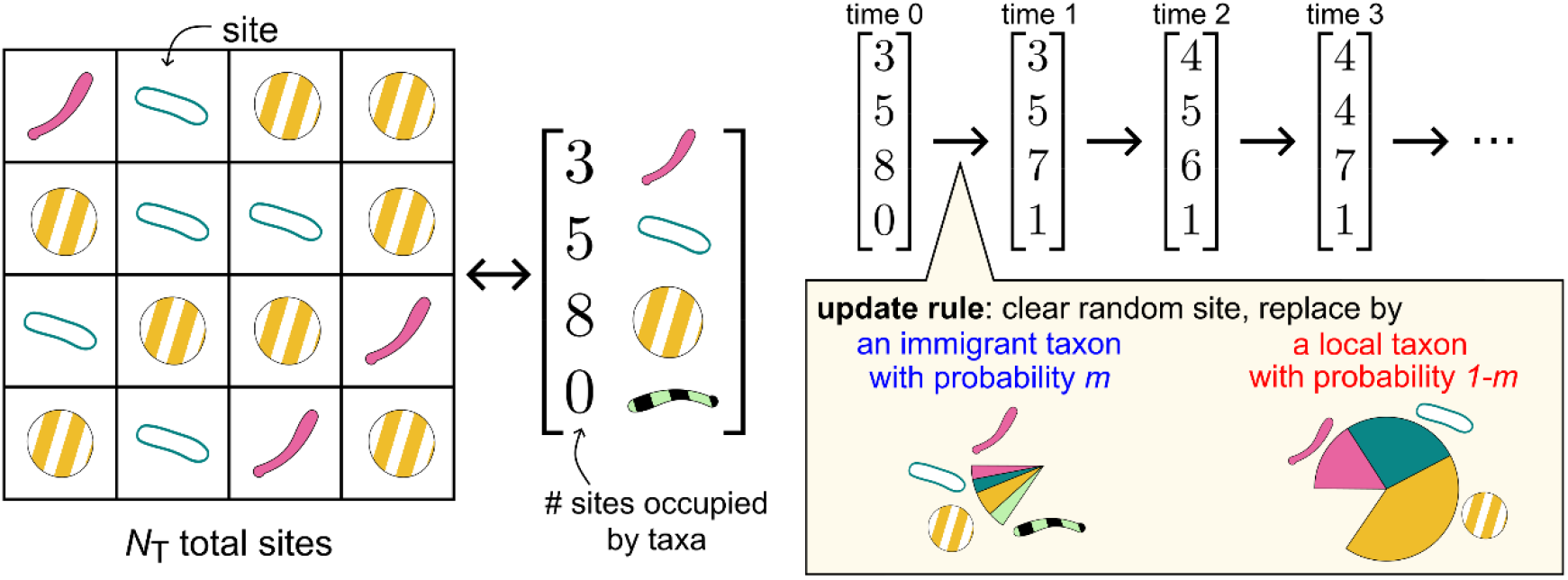
Schematic of the Neutral Community Model, describing the temporal dynamics of a ‘local’ community with *N*_T_ sites that are each occupied by some taxon *i*. At each timestep, one site is cleared and replaced by an individual from a ‘source’ community with probability *m* or by an individual from the local community with probability 1−*m*. In the source community, the relative abundance *p*_*i*_ of each taxon is fixed.

In this paper, we set out to examine how sampling affects the reliability of *N*_T_*m* inference. In any ecological sampling campaign, both the number of samples taken from a community (e.g., aliquots of biomass or count surveys) and the total number of counts in each sample (i.e., read depth in sequencing parlance) influence the accuracy and reliability of measured taxon abundances. A limited number of samples leads to an imprecise understanding of how community structure varies over space and/or time, while a limited read depth hampers the detection of rare taxa leading to an underestimation of biodiversity (12, 13). However, increasing sampling effort requires additional material and labor investment, highlighting the importance of rational sampling campaigns that optimize the allocation of resources.

In addition to sampling challenges, there are methodological limitations in how *N*_T_*m* is inferred. Though the standard ‘occupancy-based’ *N*_T_*m* inference method has been widely applied to viral (14), microbial (15–20) and macro-organismal (21, 22) datasets, it relies only on the presence/absence of each taxon, a small fraction of the total amount of information contained in abundance datasets. To complement the established occupancy-based method, here we introduce two *N*_T_*m* inference methods: a variance-based method that matches the variance in measured taxon relative abundances to the variance predicted by the NCM, and a likelihood-maximization method grounded in the Dirichlet-multinomial distribution (DM-LL). With neutral community assembly simulations, we validate these inference methods and measure their accuracy as a function of number of samples and read depth. Last, we apply these inference methods to simulated non-neutral communities and an empirical dataset to evaluate the generality of each method. Our results show that the range of immigration rates that can be inferred is limited by the read depth, and that the variance-based and DM-LL methods yield more accurate estimates than the occupancy-based method. Our findings lay out practical guidelines for quantifying immigration in complex communities using the Neutral Community Model.

## Results

### Methods to infer the immigration rate N_T_m from sampled abundance data

Following the NCM update rule (Fig. 1), the relative abundance *X*_*i*_ (of each taxon *i* out of *S* taxa) will fluctuate over time and eventually attain a stationary steady state (i.e., a statistical steady state). In the limit that the number of sites in the local community is very large, the long-term relative abundances *X*_*1*_, …, *X*_*S*_ of the local community are Dirichlet distributed (23):

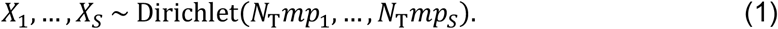

Empirical relative abundance distributions can be compared against this theoretical distribution. More pointedly, the Dirichlet distribution’s dependence on the immigration rate *N*_T_*m* means that the relative abundance distribution of each taxon is shaped by immigration, giving rise to the following variance, occupancy, and Dirichlet-Multinomial *N*_T_*m* inference methods.

### Variance-based inference

Marginalizing over the Dirichlet distribution, the relative abundance of each taxon in the local community is Beta distributed with *X*_*i*_ ∼ Beta(*N*_T_*mp*_*i*_, *N*_T_*m*(1 − *p*_*i*_)). Therefore, the mean relative abundance of a taxon in the local community is equal to its relative abundance in the source community E[*X*_*i*_] = *p*_*i*_, and its variance is inversely proportional to *N*_T_*m*, Var[*X*_*i*_] = *p*_*i*_(1 − *p*_*i*_)/(*N*_T_*m* + 1). Accordingly, *N*_T_*m* can be inferred from the observed variances and average relative abundances of each taxon. To account for sampling noise, observed variance 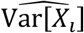 in each taxon’s relative abundance also includes a contribution from the binomial sampling variance *p*_*i*_(1 − *p*_*i*_)/*R*, where *R* is the read depth. The best fit *N*_T_*m* is obtained by least-squares regression between observed and predicted variance:

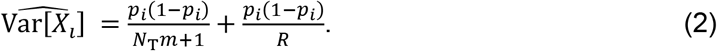

The larger the parameter *N*_T_*m*, the lower the variance of relative abundances of local-community taxa (i.e., the ‘tighter’ the relative abundance distribution across samples).

### Occupancy-based inference

A taxon’s sample occupancy *O*_*i*_ is the proportion of samples in which it has nonzero abundance (it is ‘present’). The occupancy *O*_*i*_ of neutral communities produced by the NCM can be calculated (24) by integrating the probability density function of the beta distribution from the sample detection limit *D* to 1. The detection limit is the smallest measurable value of the relative abundance, 1/*R*. The best fit *N*_T_*m* is obtained by least-squares regression between observed and predicted sample occupancies:

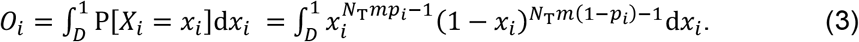

As *N*_T_*m* increases, the occupancy of all observed taxa approaches 1.

### Dirichlet-Multinomial Log-Likelihood inference (DM-LL)

Last, the Dirichlet-Multinomial Log-Likelihood (DM-LL) method identifies the *N*_T_*m* value that maximizes the likelihood of having observed the measured distribution of reads in local-community samples. Formally, this likelihood is the probability of drawing *J* samples from a multinomial distribution whose probabilities are themselves drawn from a Dirichlet distribution (Eq. 1),

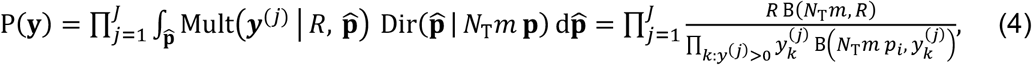

where the Dirichlet distribution captures the dependence on the relative abundance of each taxon in the source community *p*_*i*_ and *N*_T_*m*, and the multinomial distribution captures the effects of sampling. Each multinomial draw consists of *R* reads, **y**^(*j*)^ is the vector of read abundances for sample *j*, ***y*** is the set of all samples, 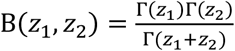 is the beta function, and 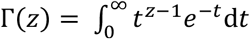 is the gamma function. The resulting Dirichlet-multinomial distribution is well-studied (25) and analytically tractable, as demonstrated by the striking simplification in Eq. 4. The maximum-likelihood estimate of *N*_T_*m*, ℒ(*N*_T_*m*|**y**), maximizes P(**y**); equivalently, it maximizes the log-likelihood:

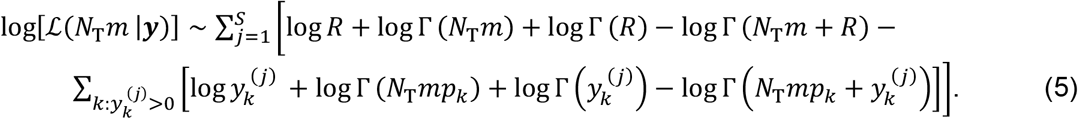

### Inference improves with increased read depth and number of samples

The Neutral Community Model posits an inverse relationship between the immigration rate and the variance of the relative abundance of each taxon Var[*X*_*i*_] in the local community according to 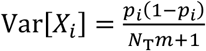. However, when communities are sampled, the estimate of a taxon’s relative abundance *X*_*i*_ itself has a variance 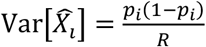 due to sampling noise, which scales inversely with the read depth *R*. When *R* < *N*_T_*m*, sampling noise dominates the ecological variance, obscuring the underlying community dynamics. Put simply, sample read depths must be larger than *N*_T_*m* for accurate inference. Since *N*_T_*m* cannot be known prior to sampling, researchers can avoid situations where inferences are limited by read depth by sequencing samples as deeply as possible (Figs. S2-S3). Increasing the number of samples improves the resolution for taxon sample occupancies, sampled abundances, and their variances.

### Variance-based and Dirichlet-Multinomial inference methods require less sampling

In this idealized sampling scenario, we explore how estimated immigration rates depend only on local-community sampling intensity and the inference method used. To this end, inferences were performed on simulated abundance datasets of neutrally assembled communities, with source-community relative abundances (Fig. S1) chosen to resemble a wastewater microbiome (Methods). Increasing the read depth and the number of samples led to more accurate and consistent *N*_T_*m* estimates (Fig. 2). Increasing the number of local-community samples increases the accuracy of all inference methods, but little improvement is achieved beyond taking 10 samples (Figs. S2 and S3). The DM-LL inference method provided the most accurate estimates of *N*_T_*m* across all combinations of read depth and number of samples, whereas the occupancy method required the largest read depth and number of samples for inference within 10% of the simulated value. Provided that a community is well described by neutral dynamics and read depths are larger than *N*_T_*m*, our findings suggest that all methods will estimate immigration rates within 10% of the true value with at least 10 local-community samples.

**Figure 2.**
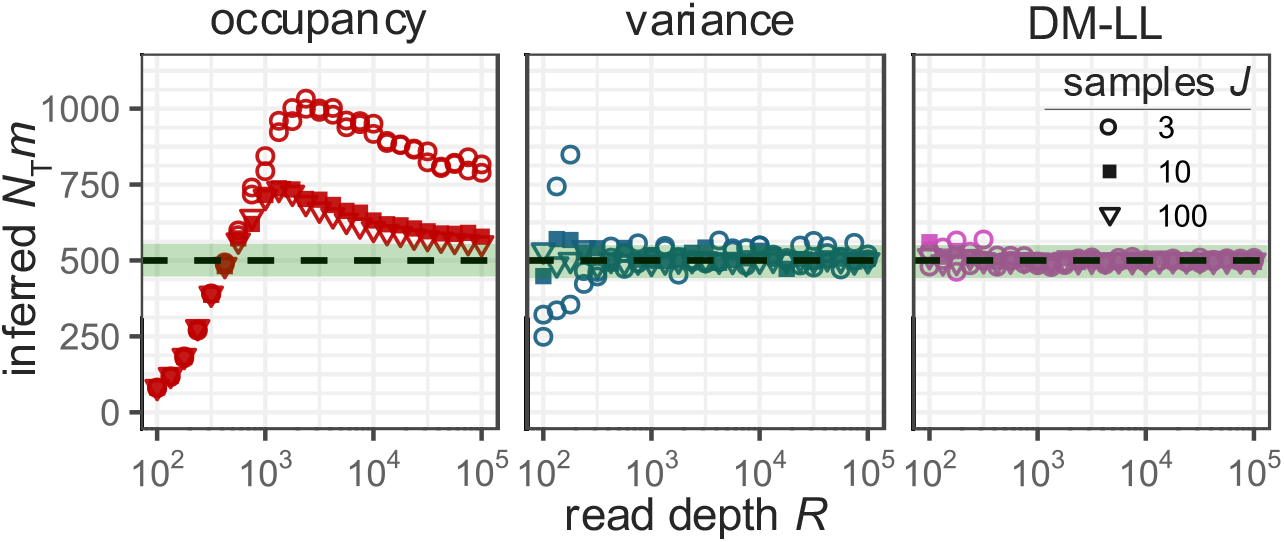
Inference accuracy depends on the sampling effort of the local community. Inferred *N*_*T*_*m* from 3, 10, and 100 local-community samples at read depths of 10^2^-10^5^ using the occupancy-based (red), variance-based (blue), and DM-LL (pink) methods. Source-community abundances were chosen to resemble a wastewater microbiome (Methods). The shaded area indicates *N*_*T*_*m* within 10% of the ground-truth value of 500 (dashed line).

### Sample a separate source community at least as much as the local community

Important ecosystems like wastewater bioreactors and gut microbiomes consist of microbes that at one point immigrated and colonized from an ‘upstream’ community (18). In the NCM, the flow of immigrants ‘downstream’ is proportional to *N*_T_*m*, and accurately estimating this quantity requires knowledge of taxa relative abundances in the upstream source community *p*_*i*_ as well as taxa relative abundances in the downstream local community. Given that the source and local communities must be sampled separately, we next investigate how sampling of the source community (which determines *p*_*i*_ in the NCM) affects *N*_T_*m* inference (Fig. 3). In the limit that sampling of the source community achieves exact estimates of *p*_*i*_, inferences achieve the accuracy of the idealized sampling scenario discussed in the previous section. Importantly, *N*_T_*m* is accurately inferred only when samples have read depths greater than *N*_T_*m*, regardless of the number of samples taken (Figs. S3). When at least 10 source-community samples are taken, typical read depths of 10^3^–10^4^ (26, 27) are sufficient to infer the immigration rate within 10% of simulated ground-truth when *N*_T_*m* ≤ 5000 (Figs. S4 and S5).

**Figure 3.**
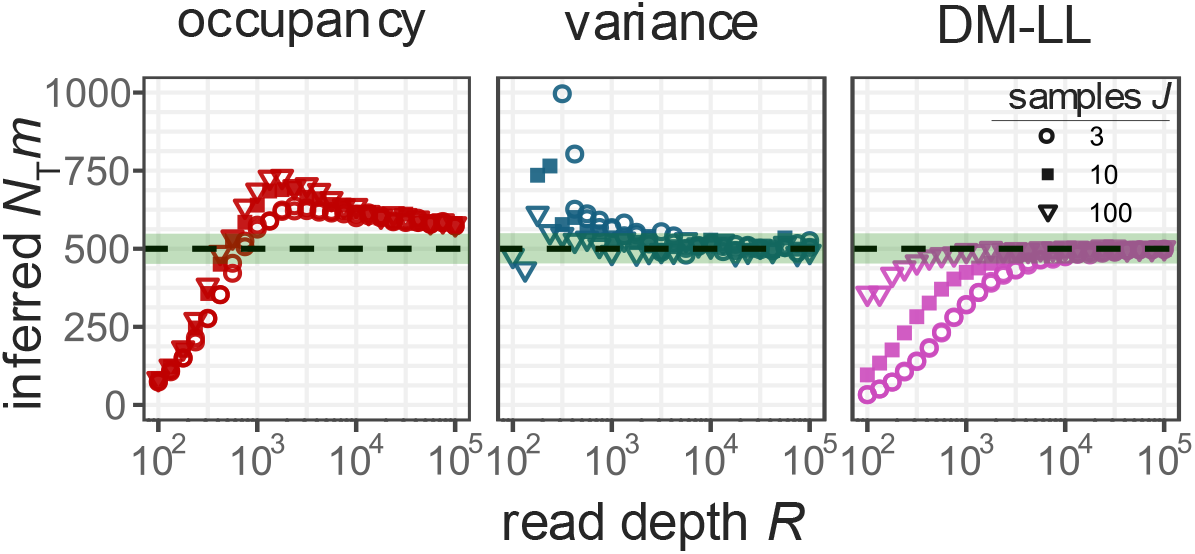
Increasing source-community sampling effort improves inference accuracy, approaching the ideal case that taxon *p*_*i*_ values are exactly known. Inferred *N*_*T*_*m* from 3, 10, and 100 source-community samples for the occupancy-based (red), variance-based (blue), and Dirichlet-multinomial log-likelihood (DM-LL, pink) methods. In all simulations, 10 samples were taken from the local community, and read depths for both source and local communities are varied from 10^2^-10^5^. The shaded area indicates *N*_*T*_*m* within 10% of the ground-truth value of 500 (dashed line).

### Quantifying immigration in non-neutral communities

In nature, communities are not strictly neutral. Selection, for example, can favor certain taxa over others, causing local-community abundances to deviate from neutral expectations. To investigate how deviations from neutrality influence *N*_T_*m* inferences, we randomly biased the relative abundance of local-community taxa by an amount proportional to a ‘non-neutrality’ parameter *σ*. Specifically, the bias *α*_*i*_ in the relative abundance of taxon *i* was drawn from a normal distribution *N(0, σ*^*2*^) where larger *σ* increases the likelihood of stronger deviations from neutrality (Methods). These biases cause some taxa to increase in abundance while others decline or disappear, perturbing the otherwise Dirichlet-distributed local community expected under purely neutral dynamics. Four values of the non-neutrality parameter *σ* were tested: 0 (neutral), 5×10^−5^ (weak), 5×10^−4^ (moderate), and 5×10^−3^ (strong). Deviations from neutrality affect the accuracy of *N*_T_*m* inference because even weak selection can drive rare taxa to extinction—eliminating any possibility of measurement—or increase their abundances far beyond neutral expectations.

The variance-based and DM-LL methods converged to accurate estimates with increasing read depth for weak and moderate levels of non-neutrality (Fig. 4 and S6). The occupancy-based method was the most sensitive to deviations from neutrality and did not converge to a stable *N*_T_*m* estimate with increasing read depth, unlike the neutral case, because previously rare taxa regularly achieved occupancies of 1 (Fig. S7). To correct this, we modified the occupancy method to only consider taxa with sample occupancies less than 1, giving better performance at weak non-neutrality (Fig. S10). Indeed, none of the inference methods could accurately infer *N*_T_*m* from local communities where strong non-neutrality impacts community structure (Figs S7-S9). At increasing levels of non-neutrality, the variance method tends to overestimate *N*_T_*m*, whereas the DM-LL and occupancy-based methods tend to underestimate *N*_T_*m*.

**Figure 4.**
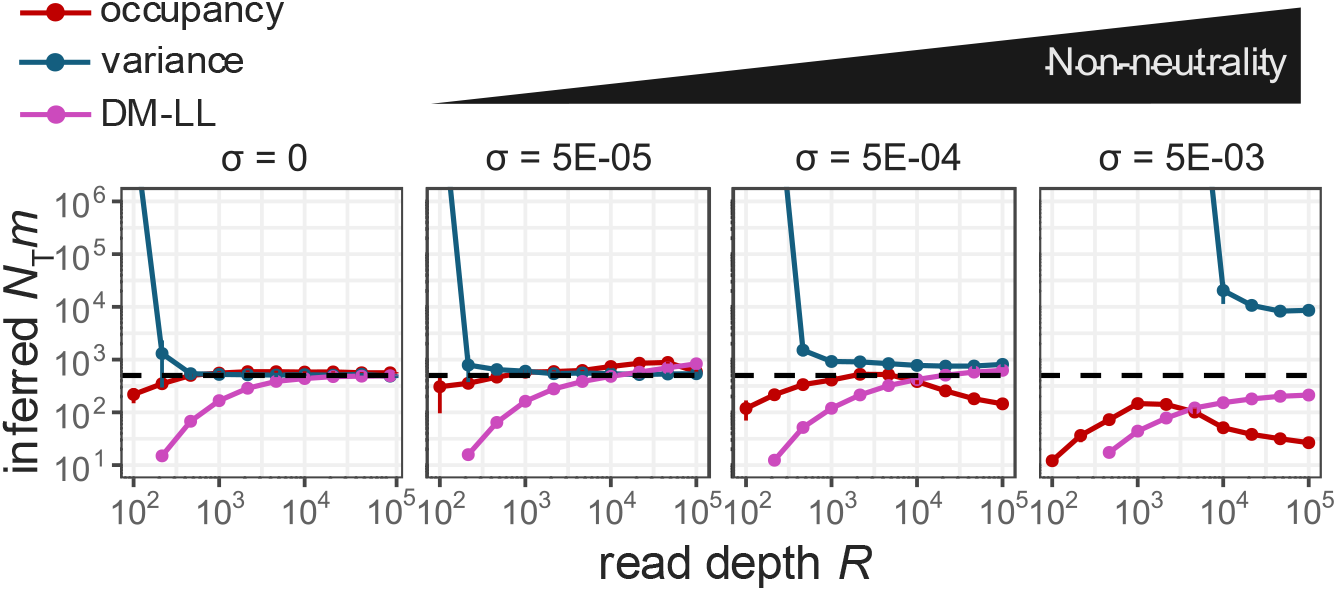
Non-neutrality reduces agreement between inference methods. Inferred *N*_*T*_*m* in neutral (*σ* = 0) and non-neutral (*σ* = 5×10^−5^, 5×10^−4^, and 5×10^−3^) local communities using the occupancy (red), variance (blue), and DM-LL (pink) methods at a simulated *N*_*T*_*m* of 500 (dashed line). Each data point consists of 10 simulations in which selection advantages are randomly drawn, local-community abundances are simulated, the source and local communities are sampled, and inference is performed. Error bars show one standard deviation.

### To infer _T_m from real data: rarefy, infer, and repeat

Finally, we apply these methods to a real 16S rRNA gene amplicon dataset consisting of samples from a full-scale activated sludge system and influent wastewater (15). This dataset provides an opportunity to validate our recommendations for sampling and *N*_T_*m* inference under natural conditions, where the true *N*_T_*m* value is unknown and additional non-neutral ecological complexities may influence community assembly. To assess inference stability and detect potential deviations from neutrality in each method, abundance tables with progressively smaller read depths were used for *N*_T_*m* inference (Methods). Inference profiles—inferred *N*_T_*m* values at varying rarefied read depths— show agreement between the estimates at read depths of 10^5^ (*N*_T_*m*_occ_ = 905, *N*_T_*m*_var_ = 1252, *N*_T_*m*_DM_ = 626) within one order of magnitude (Fig. 5); moreover, estimates from all three inference methods were stable across high read depths, confirming that inferences are not read depth limited. This approach demonstrates that rarefaction, repeated inference, and comparison across methods can provide confidence towards estimates of the immigration rate while revealing when selection or other non-neutral processes impact community assembly.

**Figure 5.**
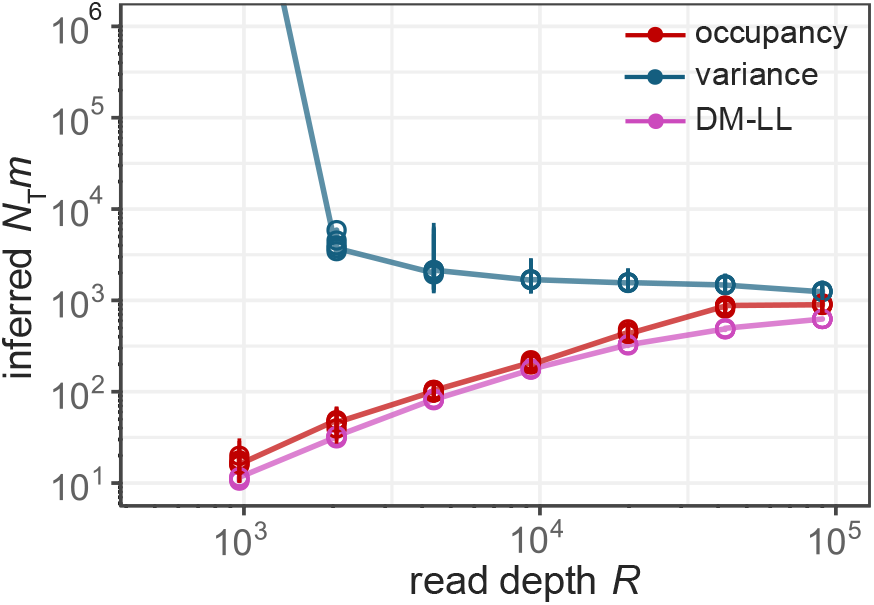
Inferring the immigration rate from wastewater to an activated sludge microbiome in a full-scale wastewater treatment plant. Inference used abundances rarefied to 5×10^2^–1×10^5^ reads per sample (Methods). Error bars show one standard deviation.

## Discussion

Pitfalls abound when attempting to quantify immigration rates using the Neutral Community Model. Motivated by our findings, we summarize key considerations here.

### What is your source community?

Originally, the NCM was formulated within the context of island biogeography (28), considering a large mainland community that serves as the source of immigrants to a smaller island community. This scale asymmetry creates a directional influence of immigration in which the island (local) community is shaped by immigration from the mainland (source) but not vice versa. On the other hand,, many ecological systems are better represented as metacommunities—networks of interacting local communities connected by dispersal without a single external source community (29). If an experimental system possesses upstream (mainland) and downstream (island) communities, both communities must be sampled separately, and the mainland-community abundances serve as the inputs for estimating the immigration rate *N*_T_*m*. By contrast, under a metacommunity framework the composition of the source community is not an independent entity but is instead the average of the local-community samples. Identifying the appropriate spatial model—mainland-island or metacommunity—impacts sampling requirements, NCM estimation, and the interpretation of estimated parameters (30, 31).

### Take at least 10 samples from the local and source communities and maximize sequence read depths

Resource constraints often limit the scale of sampling efforts, making optimized sampling strategies essential for studying community assembly. Our results show that read depth upper-bounds the *N*_T_*m* that can be inferred. Increasing the number of samples improves the precision and resolution of observed sample occupancies and improves inferences regardless of the inference method applied, although diminishing returns were observed beyond 10 local-community samples. Though more is always better, based on our simulations, we recommend taking at least 10 samples and prioritizing deep sequencing for robust *N*_T_*m* estimation. Similar recommendations apply when a source community must also be independently sampled.

### Apply all three inference methods to build confidence in estimates

The three inference methods each leverage different aspects of abundance data— occupancy, variance in relative abundances, and abundance distributions—to estimate *N*_T_*m*. Under neutral conditions, all three methods can infer *N*_T_*m* within 10% of its ground-truth value. However, non-neutrality impacts their estimates in distinct ways (Fig 4, S6). For empirical datasets, researchers should apply all three methods and evaluate the consensus between their estimates: provided that read depth is not the limiting factor, non-neutral effects likely dominate if the three methods yield inconsistent estimates. In this case, community assembly is not well described by the NCM.

### Make the most of your abundance data

In practice, read depths can vary up to 100-fold between individual samples, making normalization techniques such as rarefaction/subsampling a common preprocessing step before constructing relative abundance tables (32). When incorporated as part of a bootstrapping method, rarefaction provides a systematic way to evaluate confidence in *N*_T_*m* estimates. With evenly subsampled abundance tables, researchers can evaluate how *N*_T_*m* estimates change across read depths and determine if inferences are limited by sampling intensity or unreliable due to non-neutrality. Applying the ‘rarefy, infer, repeat’ approach in the activated sludge dataset revealed distinct patterns among the three methods across read depths. The occupancy-based, variance-based, and DM-LL methods converged to stable *N*_T_*m* estimates at higher read depths, confirming that read depths were sufficient for reliable inference. Crucially, the similarity between *N*_T_*m* estimates lends confidence to the inferred rate of microbial immigration from the influent wastewater to the activated sludge community.

### Estimating immigration rates in complex communities using the Neutral Community Model

Our results demonstrate that the NCM can address not only the qualitative question of whether community assembly is ‘neutral’ but also provide quantitative estimates of immigration rates shaping local communities. Through simulations, we codify practical recommendations for sampling and inference methodologies to guide researchers interested in applying the NCM to their study ecosystems; code that implements these inference methods is freely available. Furthermore, our ‘rarefy, infer, repeat’ approach enables researchers to extract robust immigration estimates from high-resolution sequence datasets. Although we found that *N*_T_*m* inference methods are resilient to some implementations of non-neutrality, future work should evaluate their robustness to real and simulated datasets exhibiting other forms of non-neutrality. By broadly applying these methods across diverse ecosystems and contexts, researchers can deepen ecological understanding of how dispersal influences community biodiversity.

## Materials and Methods

### Simulating neutral and non-neutral community assembly

The source-community taxa abundances were simulated by a log-series abundance distribution (Θ = 250) with a total of 10^10^ individuals and 3453 taxa (Fig. S1). Parameters for defining the source community were motivated by taxa richness in activated sludge (33), expected cell counts per litre of wastewater (34), and considerations of computational feasibility.

The neutral local-community taxa abundances were generated using a Dirichlet distribution (Eq. 1), parameterized by the source-community abundances and one of three simulated *N*_T_*m* values: 5×10^2^, 5×10^3^, and 5×10^4^. These values were chosen since pprevious studies have estimated *N*_T_*m* in this range (35–37). The NCM assumes that there are no fitness differences among taxa, so the expected relative abundances of local-community taxa mirror those in the source community.

To simulate violations of the neutrality assumption, an advantage term *α*_*i*_ is sampled from a normal distribution N(0, *σ*^*2*^) and randomly assigned for each local-community taxon *i* (20, 24, 38). The randomly drawn advantage terms have a 99.7% probability of falling within the range (−3*σ*, 3*σ*). Briefly, the *α* terms are added to Dirichlet-distributed local-community relative abundances—advantaged taxa (*α*_*i*_ *> 0*) and disadvantaged taxa (*α*_*i*_ *< 0*) have their local abundances increased and decreased relative to their source-community abundances, respectively. Taxa with relative abundances less than zero after this addition have their abundance set to zero (simulating extinction). Finally, the now ‘non-neutral’ local-community relative abundances are re-normalized to sum to one. We simulated three levels of neutrality violation (*σ* = 5×10^−5^, 5×10^−4^, and 5×10^−3^) alongside the neutral case (*σ* = 0).

All simulations, analyses, and visualizations were created using *R 4.2.2* (39).

### Sampling source- and local-community abundances

To simulate sampling and sequencing, *J* samples (each containing *R* reads) are drawn from simulated community abundances. Formally, each of these *J* samples is a draw from a multinomial distribution parameterized by the relative abundances of each taxon in the source-or local-community and by the read depth *R*.

### Simulated Scenarios

Three scenarios were used to evaluate the performance of the three *N*_T_*m* inference methods under varying ecological and sampling conditions:

- Scenario 1: Inferring _T_*m* assuming known source community abundances. ocal communities were simulated as neutral. Simulated source community abundances were used as input *p*_*i*_ for the inference methods. The number of samples taken from the local community were varied from 2 to 100. Read depths were varied from 10^2^ 10^5^.
- Scenario 2: Inferring _T_*m* in cases where source community abundances need to be sampled. ocal communities were simulated as neutral. The number of samples taken from the source community were varied from 2 to 100 to determine input *p*_*i*_ for the inference methods. Ten samples of the local community were taken. Read depths were varied from 10^2^ 10^5^.
- Scenario 3: Inferring _T_*m* from non neutral local communities. ocal communities were simulated to deviate from neutrality, testing four levels of non neutrality (σ = 0, 5×10 ^5^, 5×10 ^4^, and 5×10 ^3^). Ten samples were taken from both source and local communities. Read depths were varied from 10^2^ 10^5^.

For each combination of (i) *N*_T_*m* values, (ii) number of samples taken, and (iii) read depths, ten simulations were run; each simulation generated relative abundance datasets. The reported *N*_T_*m* is the average of the *N*_T_*m* values inferred from each simulation.

### Inferring immigration from a real wastewater bioreactor dataset

The three *N*_T_*m* inference methods were applied to 16S rRNA amplicon sequence dataset (V4 region) from a long-term sampling campaign of a full-scale activated sludge bioreactor (15). The activated sludge system and its influent wastewater were sampled twice every week over a year. The resulting dataset consists of 28 samples from each of the influent wastewater and activated sludge communities, and the read depth ranges from 31,309 to 163,733 reads per sample. Raw sequences were downloaded from the NCBI Sequence Read Archive (SRA) using the *SRA Toolkit* (https://www.ncbi.nlm.nih.gov/sra). Quality control, sequence denoising, pair-end read merging and chimera removal were performed using the *dada2* pipeline (40). Abundance tables were packaged into a *phyloseq* (41) object along with sample metadata, and used for analysis. The influent wastewater was considered the ‘source’ community, and the activated sludge system was considered the ‘local’ community. To build inference depth profiles, the abundance tables used as input for the inference methods were first rarified to different read depths before being converted to relative abundance. To ensure at least 10 samples from both source and local communities are used for inference, the maximum read depth considered was chosen to be 10^5^.

## Data, material and software availability

R code that replicates Figs. 2-5 and that can be applied to any sequencing dataset is available at https://github.com/ramisrafay/NCM-Inference-Methods

## Acknowledgments

This work was supported by an SFU Graduate Deans Entrance Scholarship awarded to R. R., Banting and Pacific Institute for the Mathematical Sciences Postdoctoral Fellowships awarded to E.W.J., a Natural Sciences and Engineering Research Council of Canada (NSERC) Discovery Grant and Discovery Accelerator Supplement RGPIN-2020-04950 and a Tier-II Canada Research Chair CRC-2020-00098 awarded to D.A.S., and an NSERC Discovery Grant RGPIN-2020-05086 awarded to S.J.F.

