## Supplementary Figures for "How to quantify immigration from community abundance data using the Neutral Community Model"

Ramis Rafay[a], Eric W. Jones[b,c,d], David A. Sivak[d], S. Jane Fowler[a]
[a]*Department of Biological Sciences, Simon Fraser University, Burnaby, BC, Canada V5A1S6*
[b]*Department of Physics and Energy Science, University of Colorado Colorado Springs, Colorado Springs, CO, USA 80918*
[c]*BioFrontiers Center, University of Colorado Colorado Springs, Colorado Springs, CO, USA 80918*
[d]*Department of Physics, Simon Fraser University, Burnaby, BC, Canada V5A1S6*

**This PDF file includes:**

Supplementary Figures S1-S10

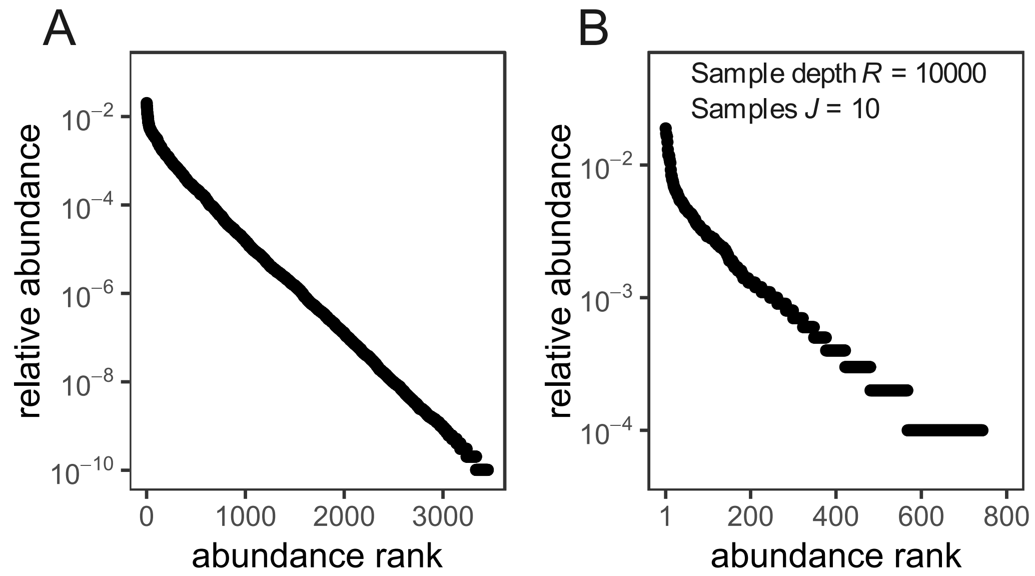


Figure S1. (A) The 'true' source-community rank abundance distribution is drawn from a logseries distribution with parameters N = $10^{10}$ and alpha = 250. (B) An example of the 'sampled' source-community rank abundance distribution produced by sampling the true source-community abundances using 10 samples with 10,000 reads each.

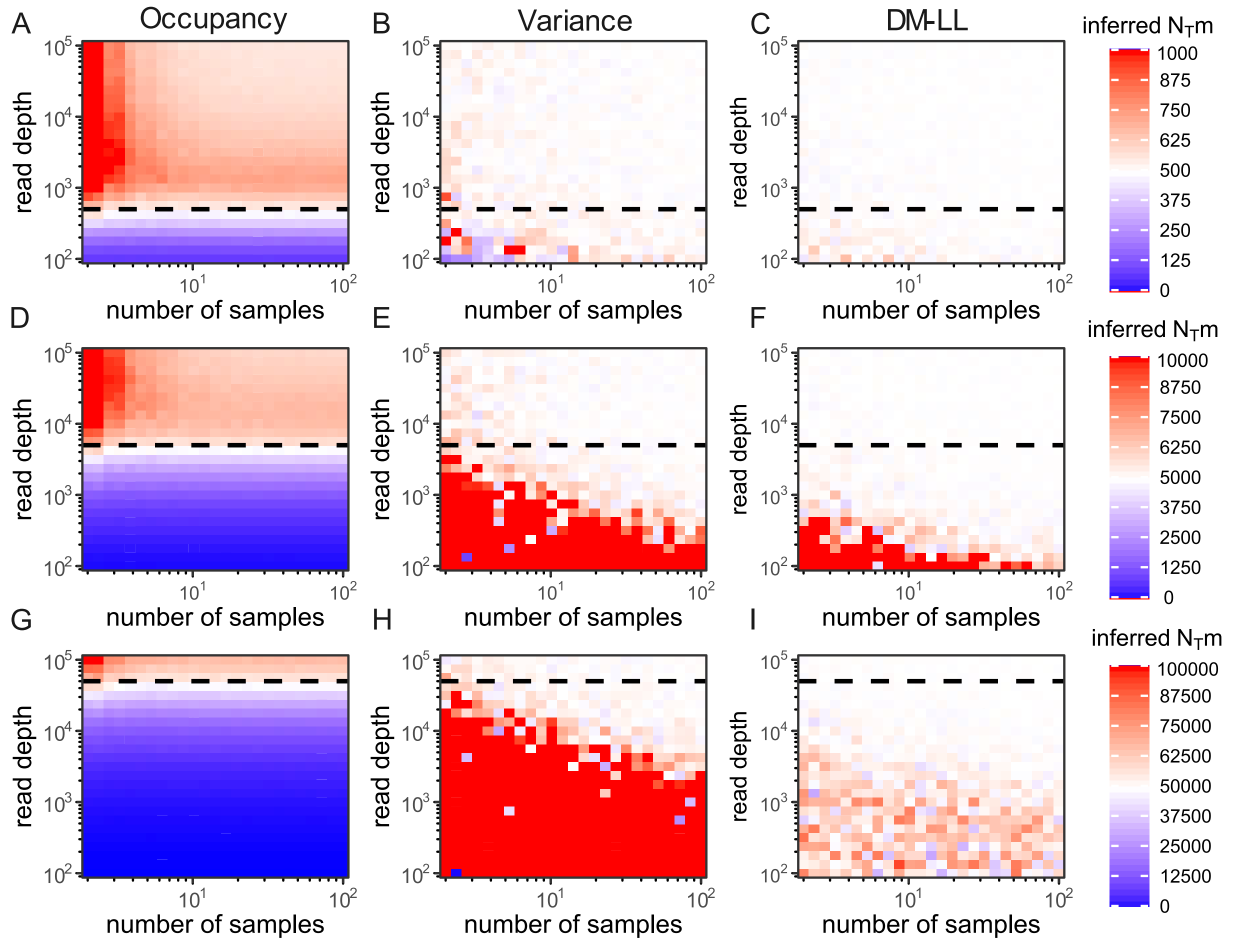


Figure S2. Heatmaps showing inferred $N_T m$ using the occupancy-based, variance-based, and DM-LL methods across varying numbers of local-community sample replicates and read depths. Simulations of community assembly, sampling and inference were performed 10 times each for every combination of number of local-community samples (varied from 2-100) and read depths (varied from $10^2$-$10^5$ reads per sample). Dashed lines indicate simulated $N_T m$: $5\times10^2$ (A-C), $5\times10^3$, (D-F), $5\times10^4$ (G-I). The color gradient depicts whether the inference method underestimated (blue), correctly estimated (white) or overestimated (red) $N_T m$.

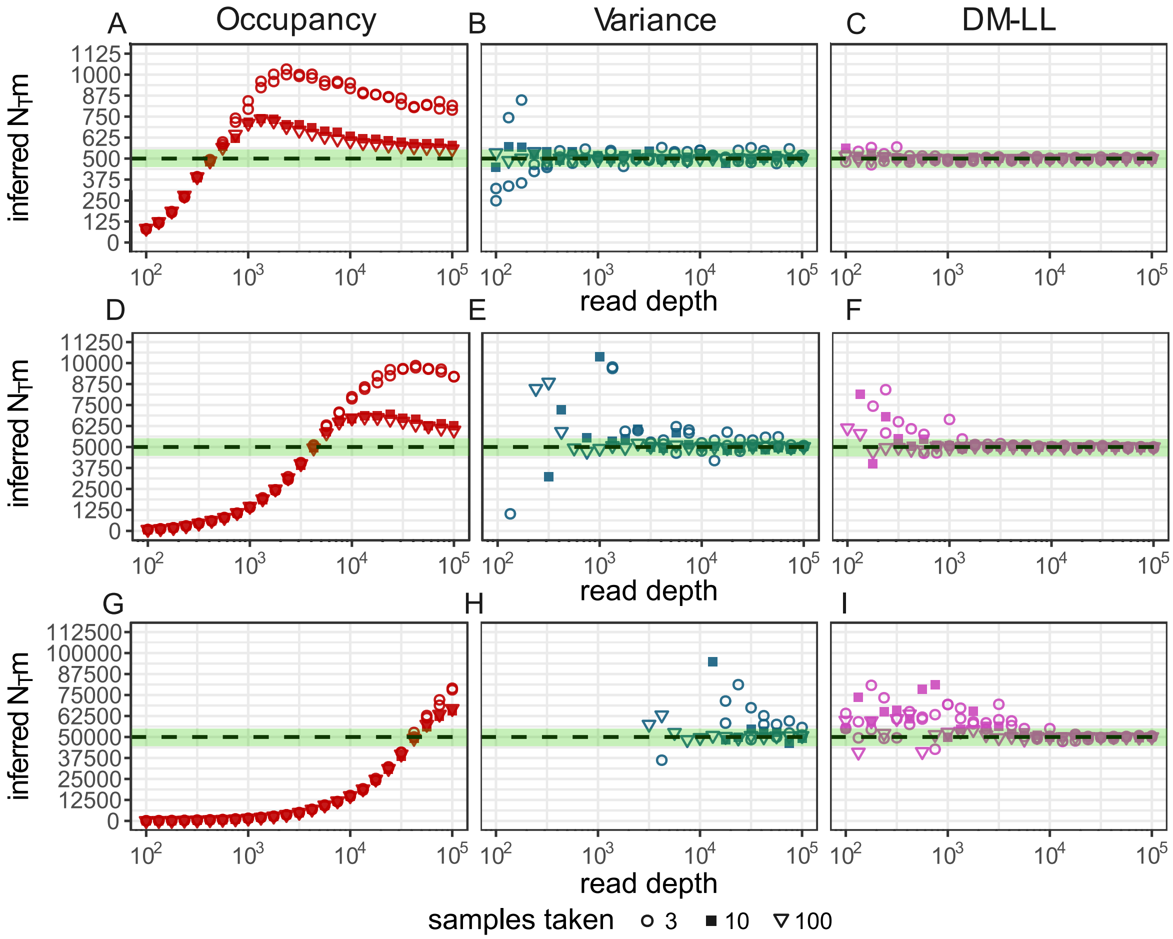


Figure S3. Accurate, reliable inferences require read depths of local-community samples larger than $N_T m$. Inferred $N_T m$ from 3, 10, and 100 local-community samples at read depths of $10^2$-$10^5$ using the occupancy-based, variance-based, and DM-LL methods. Dashed lines indicate simulated $N_T m$: $5 \times 10^2$ (A-C), $5 \times 10^3$, (D-F), $5 \times 10^4$ (G-I). All methods require high read depth to accurately infer large $N_T m$, with inference accuracy improving as read depth increases.

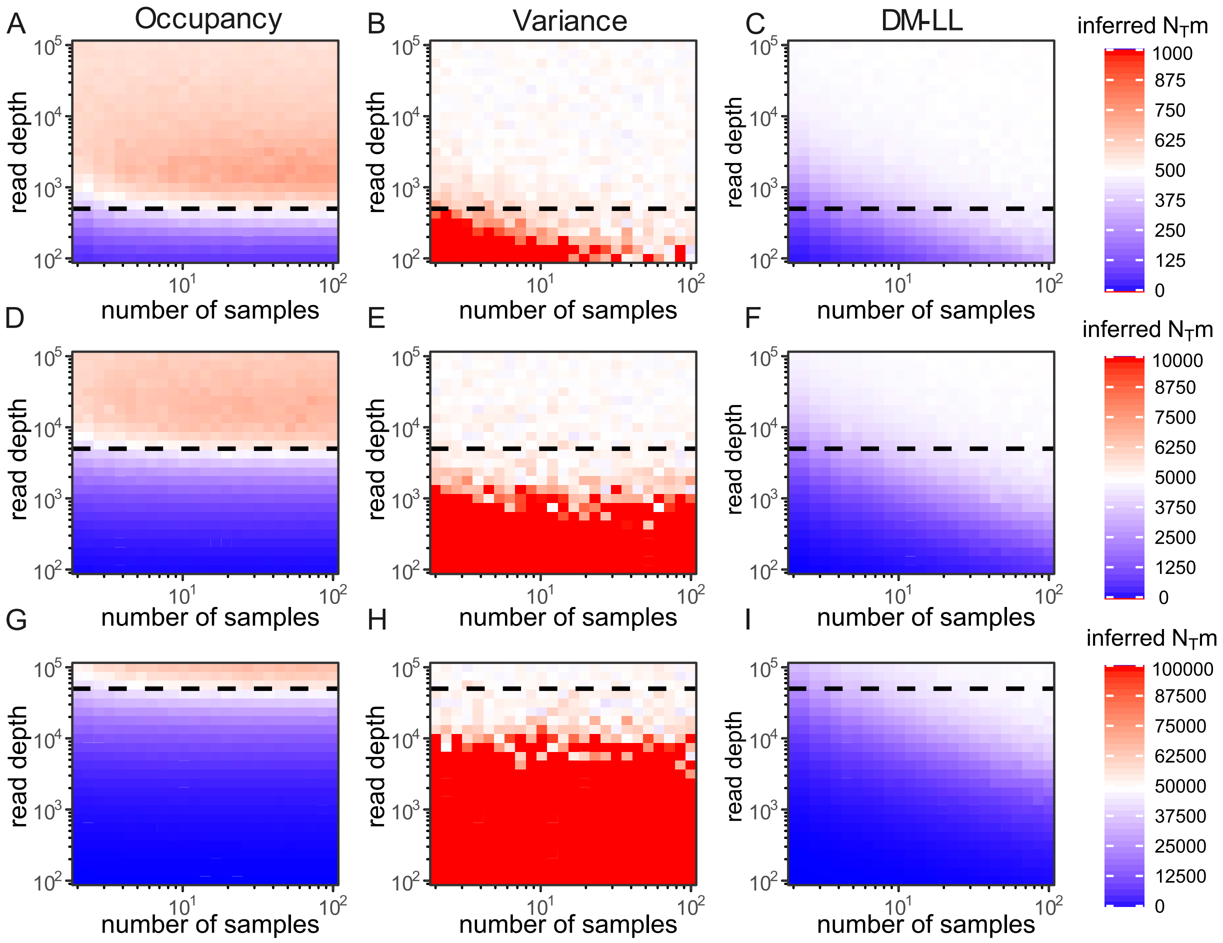


Figure S4. Same as in Fig. S2, except here the source community is sampled, leading to noisy estimations of $p_i$ that are used for $N_T m$ inference. In each case, 10 samples were taken from the local community. Dashed lines indicate simulated $N_T m$: $5{\times}10^2$ (A-C), $5{\times}10^3$, (D-F), $5{\times}10^4$ (G-I). The color gradient depicts whether the inference method underestimated (blue), correctly estimated (white) or overestimated (red) $N_T m$.

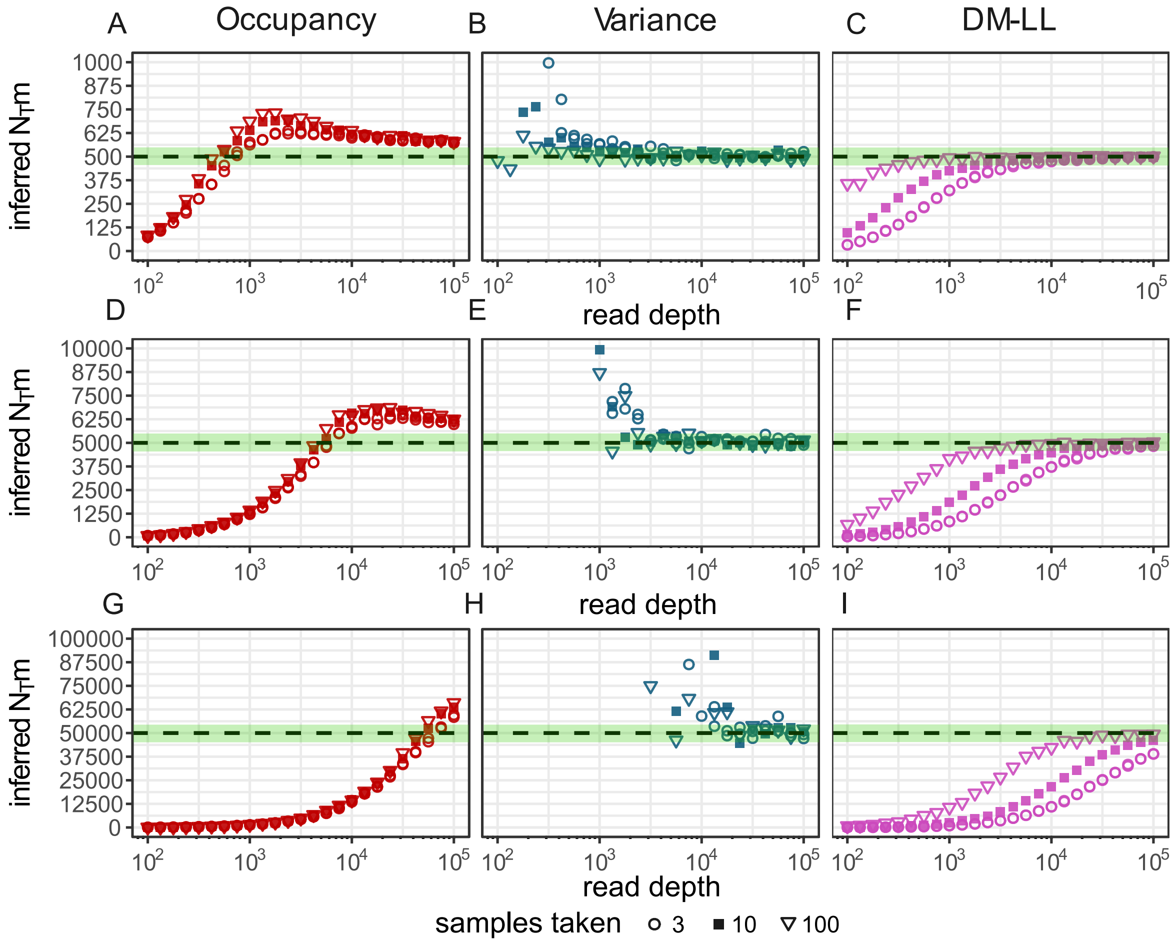


Figure S5. Same as in Fig S3, except here the source community is sampled, leading to noisy values of $p_i$ that are used for $N_T m$ inference. Dashed lines indicate simulated $N_T m$: $5\times10^2$ (A-C), $5\times10^3$, (D-F), $5\times10^4$ (G-I).

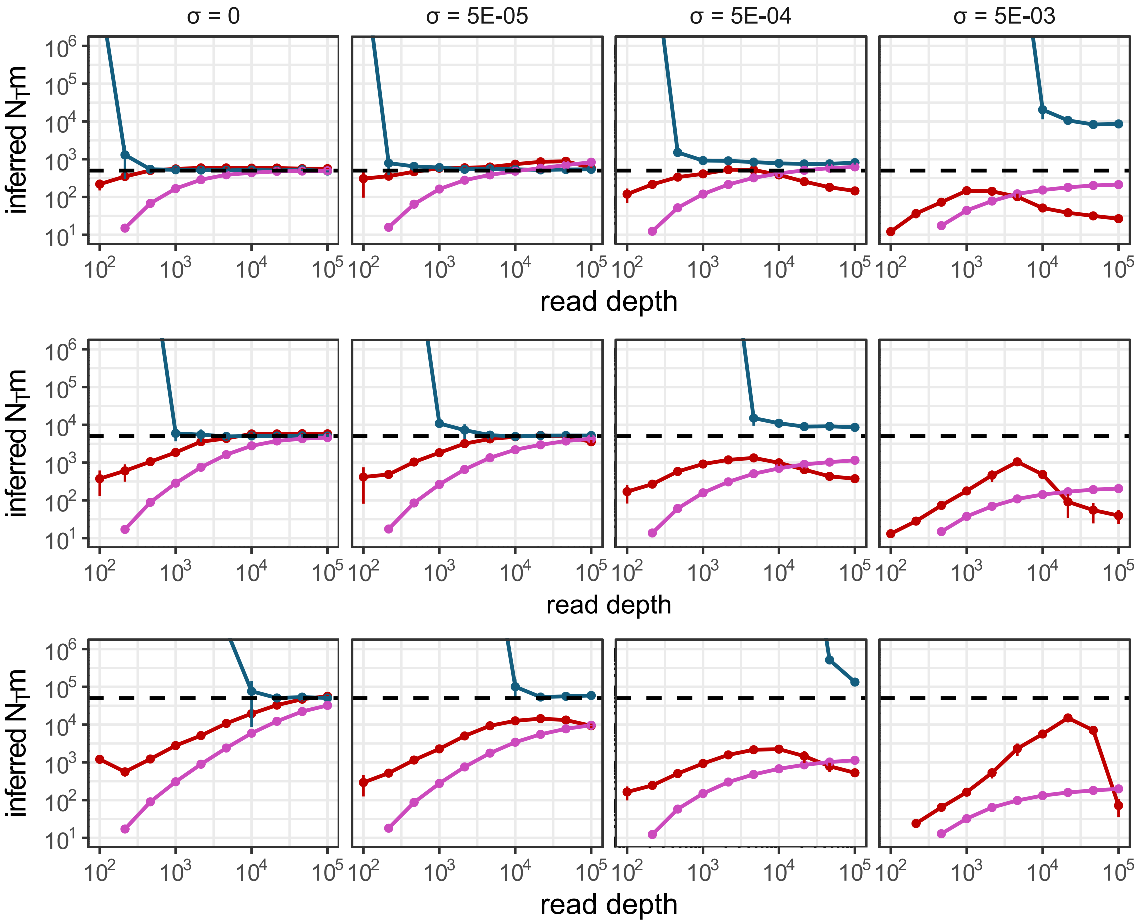


Figure S6. $N_Tm$ inference under non-neutrality (as in Fig. 4) for three different ground-truth $N_Tm$ values: $5\times10^2$ (top), $5\times10^3$, (middle), $5\times10^4$ (bottom). Inferred $N_Tm$ from the occupancy-based (red), variance-based (blue), and DM-LL (pink) methods at levels of non-neutrality parameter $\sigma$: 0 (neutral), $5\text{x}10^{-5}$ (weak), $5\text{x}10^{-4}$ (moderate), and $5\text{x}10^{-3}$ (strong). Dashed lines indicate simulated $N_Tm$.

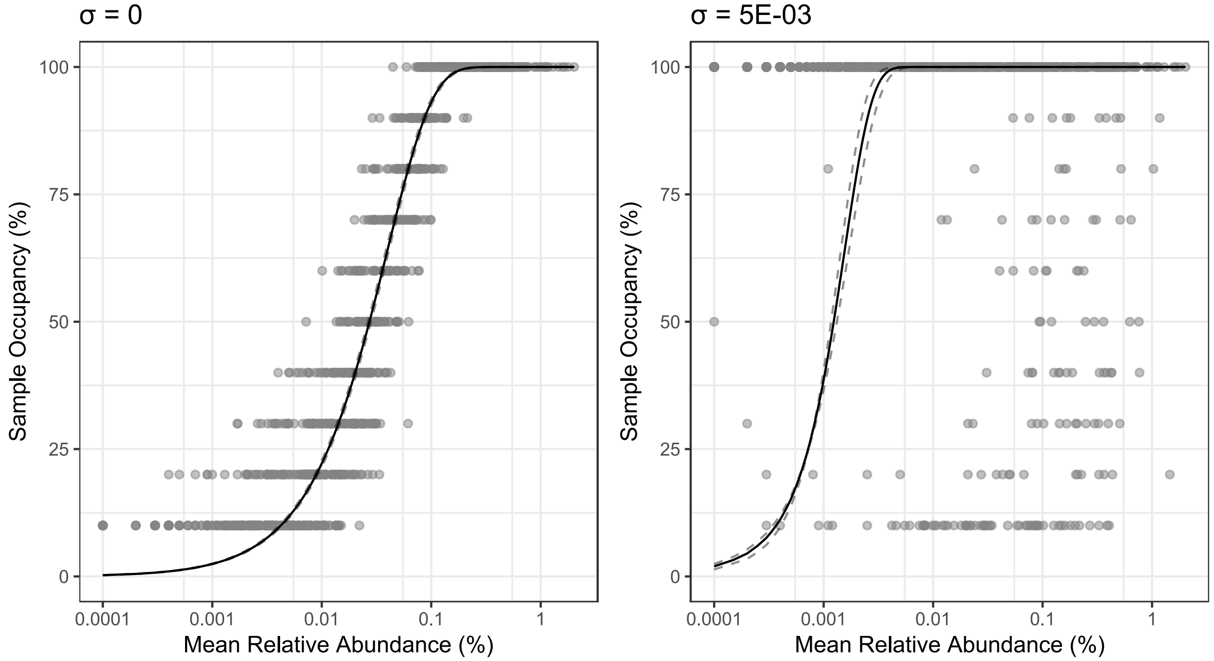


Figure S7. Diagnostic plot for the occupancy-based inference method, indicating the fit of the model (curve) to observed sample occupancies (points). Model fit expected under ideal neutral assembly (left) is disrupted by strong non-neutrality (right), leading to incorrect $N_T m$ estimations.

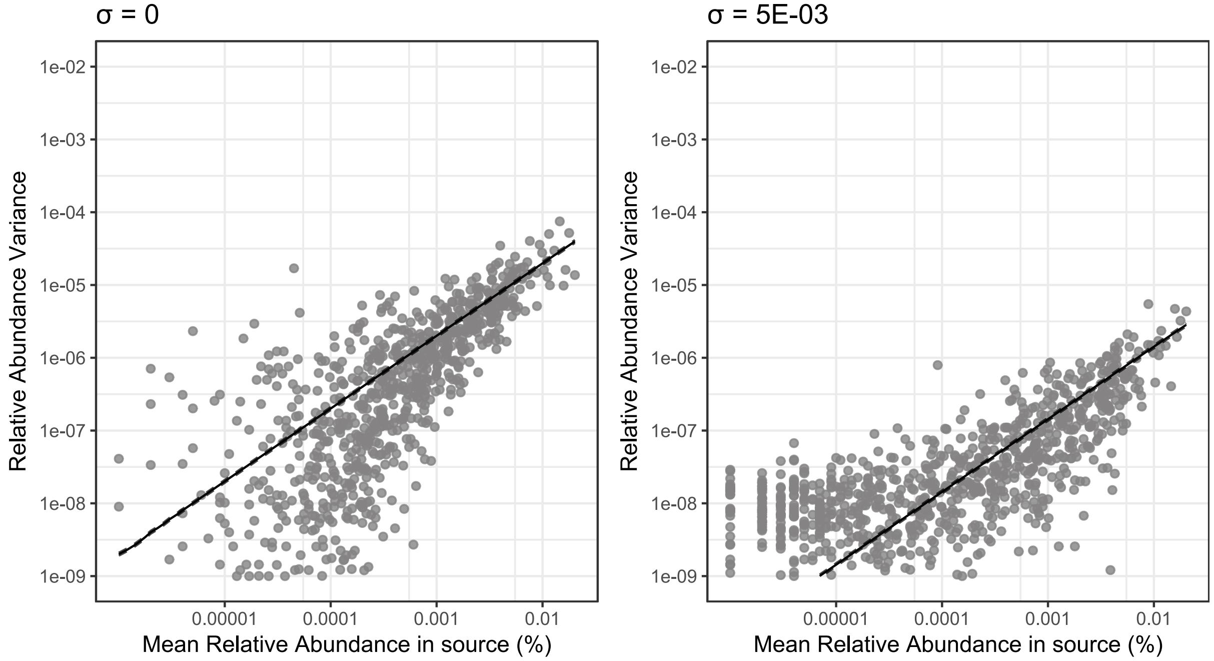


Figure S8 Diagnostic plot for the variance-based inference method, indicating the fit of the model (line) to observed taxon relative abundance variances (points). Model fit expected under ideal neutral assembly (left) is disrupted by strong non-neutrality (right), leading to incorrect $N_T m$ estimations.

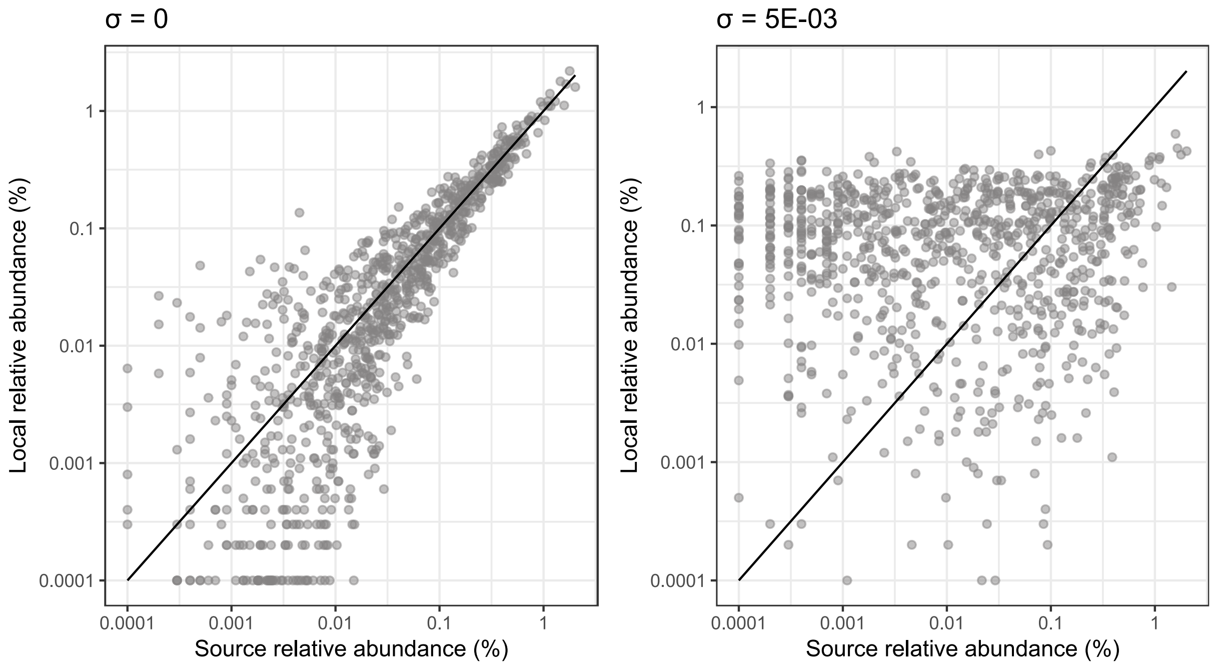


Figure S9. Diagnostic plot for the Dirichlet-multinomial inference method. The expected relationship between source- and local-community relative abundances—that they are equal on average—under neutrality (left) is disrupted by strong non-neutrality (right), leading to incorrect $N_T m$ estimations.

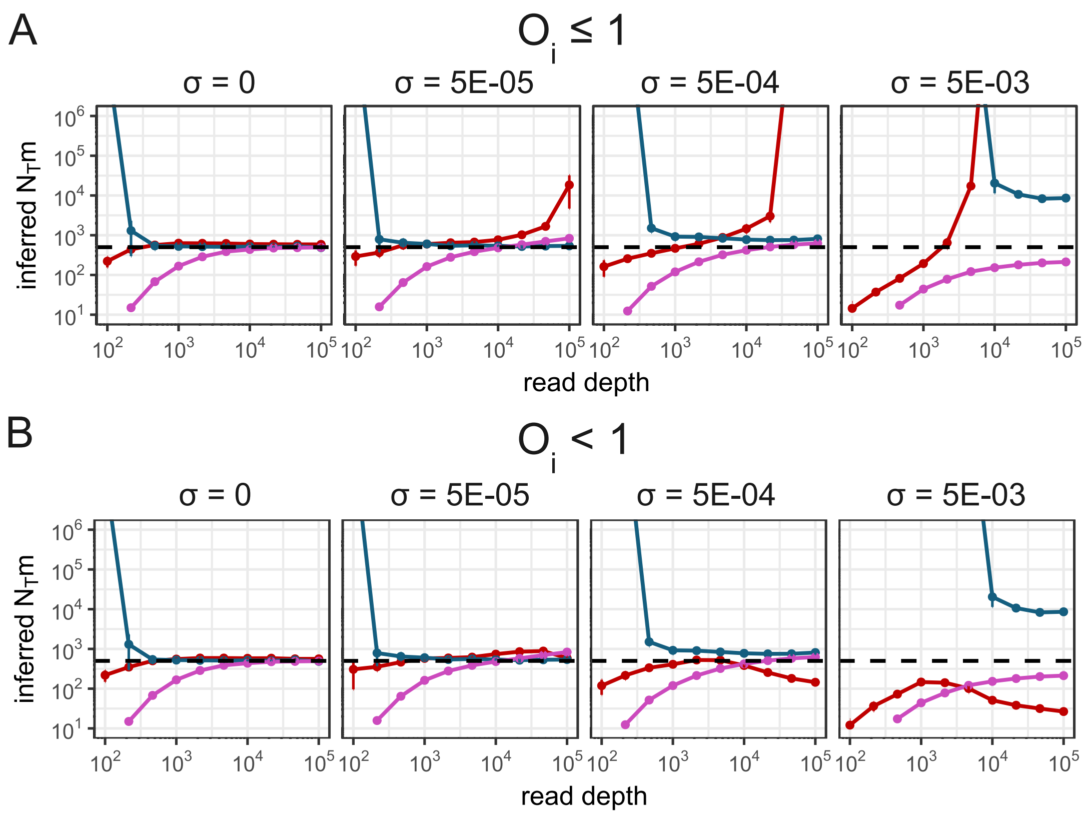


Figure S10. The occupancy-based inference method is particularly sensitive to non-neutrality effects if taxa with sample occupancies of 1 are included in estimation (A). Simply excluding species with occupancies of 1 improves occupancy-based inference from non-neutral local communities and has no adverse impact on performance in the neutral case (B).
